# Brain intrinsic connection patterns underlying tool processing in human adults are present in neonates and not in macaques

**DOI:** 10.1101/2021.12.17.473098

**Authors:** Haojie Wen, Ting Xu, Xiaoying Wang, Xi Yu, Yanchao Bi

## Abstract

Tool understanding and use are supported by a dedicated left-lateralized, intrinsically connected network in the human adult brain. To examine this network’s phylogenic and ontogenetic origins, we compared resting-state functional connectivity (rsFC) among regions subserving tool processing in human adults to rsFC among homologous regions in human neonates and macaque monkeys (adolescent and mature). These homologous regions formed an intrinsic network in human neonates, but not in macaques. Network topological patterns were highly similar between human adults and neonates, and significantly less so between humans and macaques. The premotor-parietal rsFC had most significant contribution to the formation of the neonate tool network. These results suggest that an intrinsic brain network potentially supporting tool processing exists in the human brain prior to individual tool use experiences, and that the premotor-parietal functional connection in particular offers a brain basis for complex tool behaviors specific to humans.

## Introduction

Complex and flexible tool making and tool use are argued to be unique to homo sapiens (Ambrose, 2001; Gibson et al., 1994; Oakley, 1956; Vaesen, 2012; Laland and Seed, 2021). Such abilities are aligned with observations of a dedicated left-lateralized network in human adult brain that is particularly relevant for processing tools (e.g., hammers, axes, scissors), which includes left lateral occipital-temporal cortex (LOTC), inferior and superior parietal lobule, inferior frontal gyrus, and premotor cortex. These regions are preferentially activated when human adults view pictures of tools relative to other types of objects (e.g., faces, animals, and large, non-manipulable objects), listen to tool names, imagine tool use, or pantomime tool use (Chao and Martin, 2000; Chouinard and Goodale, 2012; Lewis, 2006; Peelen et al., 2013; Wang et al., 2018). Also, they have been observed to be both structurally and functionally connected (Bi et al., 2015; Konkle and Caramazza, 2017; Peelen et al., 2013) and lesions to these regions and/or to their underlying white matter connections can lead to deficits in tool understanding and use (Bi et al., 2015; Buxbaum et al., 2014; Garcea et al., 2020; Tarhan et al., 2015). It is assumed that each of these regions contributes distinct computations to tool processing, with the LOTC, as part of the ventral visual pathway, involved in visual shape analysis and, optimally connected to the parietal and frontal regions to support grasping and manipulating tools (Mahon, 2020).

What are the phylogenic and ontogenetic origins of the tool processing network observed in human adults? Is its formation simply the result of associative learning, based on individual experiences of tool manipulation, which bridge sensory and motor representations of tools? Or is this network (partly) innate, predisposed in the human brains prior to any individual object use experience, and potentially human-specific, given its evolutionary significance in homo sapiens? One way to tackle this fundamental question is to compare this brain system in human adults with that in human neonates and nonhuman animals. However, it is difficult to perform tool-processing experimental tasks with human neonates not only for practical reasons but also because of highly limited cognitive/motor skills (only reflective motor responses). The approach we took here, motivated by the notion that brain function is determined by connection patterns (Passingham et al, 2002), is to take advantage of resting-state functional connectivity (rsFC) data and examine whether the intrinsic brain connection pattern among the homologous brain regions of interest are already in place in human neonates.

Regarding nonhuman primates, similarities and differences to humans have been reported on both behavioral and neural levels. The simpler forms of tool use and even tool making are not unique to humans. Animals have visual and motor experiences with objects such as gasping a stick, and some have even demonstrated simple tool use (e.g., apes and crows use sticks to forage for insects, Bentley-Condit and Smith, 2010; Fayet et al., 2020; Shumaker et al., 2011). Nevertheless, humans are arguably the only species that can make and use sophisticated tools based on causal (mechanical) understanding of the relationship between its physical properties, use and function (Johnson-Frey, 2003; Laland and Seed, 2021; Osiurak and Reynaud, 2020; Penn et al., 2008; Vaesen, 2012; Visalberghi and Limongelli, 1994), which allows them to convert ordinary objects into tools for flexible functional use as early as two years of age (Kastner et al., 2017). Neurally, in the macaque brains, object grasping or simple-tool-use is supported by a lateral network encompassing the parietal cortex, premotor area and inferior frontal gyrus (e.g., Borra et al., 2017; Obayashi et al., 2001), regions homologous or in close proximity to the tool-processing areas in human adults; Yet, species-differences were observed in the left inferior parietal cortex associating with tool-use-activities (Peeters et al., 2009). The characteristics of brain connectivity pattern among these homologous brain regions, however, have not been compared across species and developmental trajectories.

Here, we empirically tested the phylogenic and ontogenetic origins of the intrinsic tool processing network observed in human adults by comparing resting-state functional connectivity (rsFC) pattern among tool processing regions in human adults (n=100) to rsFC among homologous regions in human neonates without any motor (n = 118) and in macaque monkeys with ample visual/motor experiences (adolescent and mature, n = 25). If the emergence of the intrinsically connected tool-processing network in human adults is driven by learnt sensory-motor association based on the visual-motor/manipulation experiences with objects, then similar intrinsic connectivity pattern among the homologous regions are not predicted in human neonates, who have not developed any nonreflective motor skills and thus no object use experience, but predicted in macaques, who have extensive motor experience with objects. Alternatively, the tool network observed in human adults may be innate and (at least partly) unique to homo sapiens, supporting the human-unique complex tool making/use behaviors, and we would expect to observe similar intrinsic connectivity pattern already present among the homologous regions in human neonate brain, and not in the macaque brain. A brain network supporting face processing (henceforth face processing network), which has been reported for both humans (Wang et al., 2016) and macaques (Schwiedrzik et al., 2015), was also assessed in these three populations as a reference point for the potential tool processing network.

## Results

### Intrinsic functional connectivity results

#### Human adult tool network characterization

Nodes (regions-of-interest; ROIs) of the neural networks underlying tool (and face as a control domain) processing in human adults were objectively generated from meta-analyses based on the Neurosynth database incorporating 14,371 fMRI studies in total (https://neurosynth.org, version 0.7 released July, 2018, Yarkoni et al., 2011). The tool processing network, derived from 115 studies using the term “tools,” contained four regions in the left hemisphere: left lateral occipitotemporal cortex (LOTC), left inferior frontal gyrus (LIFG), left premotor gyrus (LPreG), and left inferior and superior parietal lobule (LIPL/SPL). The face processing network was derived from 896 studies (term “face”) and constituted eight regions: bilateral fusiform face areas (LFFA and RFFA), occipital face areas (LOFA and ROFA), superior temporal gyri (LSTG and RSTG), right inferior frontal gyrus (RIFG) and right anterior temporal lobe (RATL, Figure 1A). Using the resting-state dataset in the Human Connectome Project (HCP, Van Essen et al., 2013, n = 100, randomly selected, mean age = 28.3 ± 3.4), we demonstrated that these regions being consistently activated by tools (or faces) constituted tightly connected networks, replicating previous literature (Peelen et al., 2013; Stevens et al., 2015; Wang et al., 2016). That is, the rsFC among the four tool processing nodes (i.e., six within-tool-domain connections) and among the eight face processing nodes (i.e., 28 within-face-domain connections) were greater than the rsFC between tool and face processing nodes (i.e., 32 between-domain connections; see Figure 1B for an analysis schema; within-tool-domain > between-domain: *t*99 = 15.4, *p*corrected < 0.001, *Cohen’s d =* 1.54; within-face-domain > between-domain: *t*_99_ = 15.4, *p*_corrected_ < 0.001, *Cohen’s d =* 1.54, Figure 2A).

**Figure 1.**
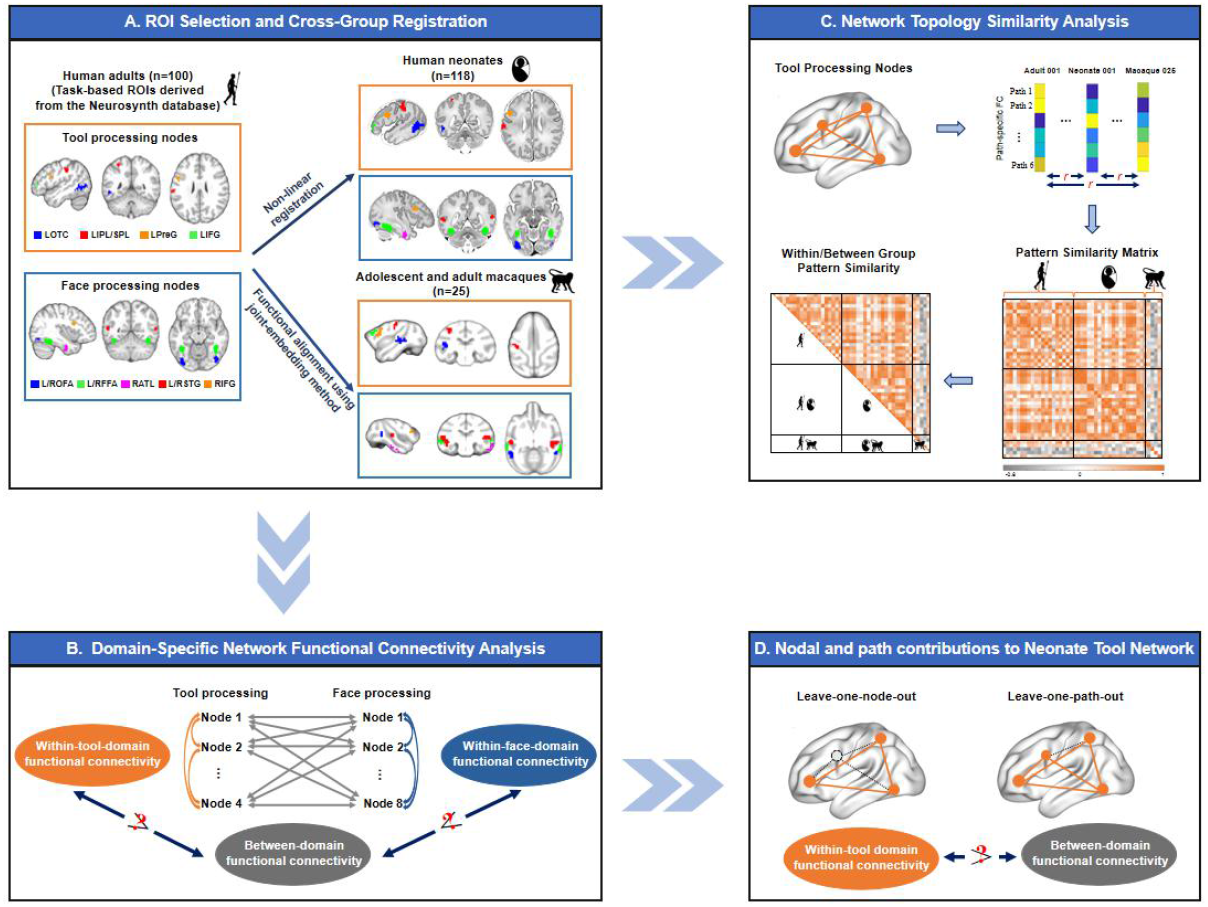
Flow chart of resting-state functional connectivity (rsFC) analyses for tool and face processing networks in human adults, human neonates and macaques. A. Tool and face processing nodes are presented in slice views on standard templates for human adults (T1-weighted), human neonates (T2-weighted), and macaques (T1-weighted). These nodes were initially derived from a task-based fMRI meta-analysis using the Neurosynth database and then registered to human neonate and macaque spaces using non-linear registration and functional alignment approaches, respectively. B. Intrinsic networks were first evaluated by comparing rsFC between nodes of the same network to that of nodes belonging to different networks. C. A step-by-step procedure is illustrated for computing network topology similarity between groups (using the tool processing network as an example). D. Additional characterization of nodal and path contributions to the formation of the tool processing network in human neonates using the leave-one-out approach. Slice views and projected brain images were prepared in Mricron (https://www.nitrc.org/projects/mricron) and BrainNet Viewer (Xia et al., 2013), respectively. LOTC: left lateral occipitotemporal cortex; LIPL/SPL: left inferior and superior parietal lobule; LPreG: left premotor gyrus; LIFG: left inferior frontal gyrus; L/ROFA: left and right occipital face areas; L/RFFA: left and right fusiform face areas; RATL: right anterior temporal lobe; L/RSTG: left and right superior temporal gyri; RIFG: right inferior frontal gyrus.

**Figure 2.**
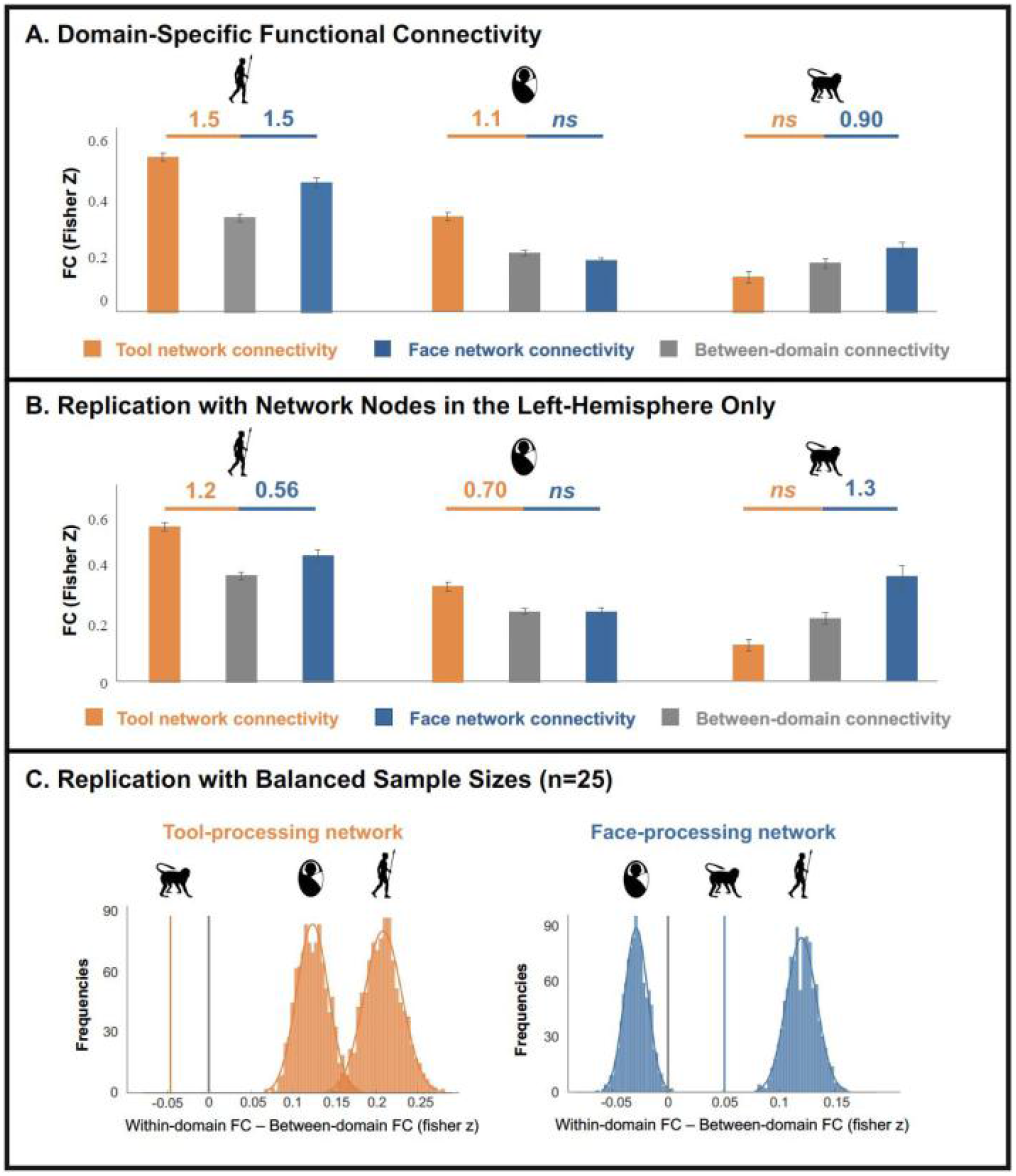
The human adult tool-network intrinsic connectivity structure is present in human neonates, but not in adolescent/mature macaques. A. Bar graphs illustrate resting-state functional connectivity (rsFC) values for within-domain and between-domain connectivity in all three groups. Effect sizes (Cohen’s *d*) are shown for comparisons in which significantly greater rsFC than between-domain rsFC was observed (all *p*_corrected_ < 0.001). Error bars indicate standard errors. B. Bar graphs depict replication results for the network analysis using only left-hemispheric nodes with balanced within-tool-domain and between-domain nodal distance. Effect sizes (Cohen’s *d*) are shown for comparisons in which significantly greater rsFC than between-domain rsFC was observed (all *p*_corrected_ < 0.001). Error bars indicate standard errors. C. Distribution maps show network effects from the bootstrapping analysis (n=10,000) in human adults and neonates, which were calculated as within-domain minus between-domain rsFC (rsFC differences) for tool and face processing networks. A single line is used to indicate the rsFC differences for macaques, since no bootstrapping analyses were performed in this group. The grey line represents 0.

#### Tool homologous intrinsic network structure present in human neonates

Tool and face processing nodes identified above in human adults in MNI152 space were transformed onto 40-week templates available on the developmental human connectome project (dHCP) website (https://gin.g-node.org/BioMedIA/dhcpvolumetric-atlas-groupwise), using Advanced Normalization Tools (ANTs, Avants et al., 2009, https://stnava.github.io/ANTs, Figure 1A). Hereafter, they are referred to as tool (or face) homologous nodes or regions. Network functional connectivity strengths were calculated using neonate resting state data (n = 118) available from the dHCP (see inclusion criteria in the methods). An intrinsic functional network among the tool homologous regions was observed. That is, within-domain rsFC for the tool homologous network was significantly greater than between-domain rsFC (*t*_117_ = 12.4, *p*_corrected_ < 0.001, *Cohen’s d =* 1.1, Figure 2A). The same results held when the pre-term and full-term neonates were analyzed separately (full-term neonates: *t*_105_ = 11.4, *p*_corrected_ < 0.001, *Cohen’s d =* 1.1; pre-term neonates: *t*_11_ = 6.3, *p*_corrected_ < 0.001, *Cohen’s d =* 1.8, Figure S1A). No coherent face homologous network was observed in neonates, as within-face-domain rsFC was not stronger than between-domain rsFC (*t*_117_ < 0).

#### Tool homologous intrinsic network structure absent in macaques

The tool and face processing nodes identified in human adults were registered to the macaque 112RM-SL templates (McLaren et al., 2009, 2010) through a combination of transformations generated by alignment methods recently developed for cross-species transformation (Xu et al., 2020) and ANTs (Figure 1A, see Methods for the details of the cross-species registration). Given the challenge of cross-species brain registration, we took further care to check the transformed ROIs in the macaque brain (Figure 1A) in relation to potentially tool-relevant landmarks reported in the literature. The registered frontal and parietal ROIs corresponded largely to the key components of the macaque grasping network (Borra et al., 2017), including the F5 (for PreG), m46v/m12r (for LIFG) and AIP (anterior intraparietal, for LIPL/SPL). Specifically, they were overlapping with these regions manually drawn and available in Howells et al. (2020, Figure S2). The transformed LOTC node lays at the intersections of the occipital and the temporal lobes of the macaque brain. Finally, using the same registration method, ROIs homologous to the nodes of the face network (control) were also generated in the macaque brain with locations largely consistent with task-based activations in macaques (Hesse and Tsao, 2020; Ku et al., 2011; Landi and Freiwald, 2017; Tsao et al., 2008). The open-source resting-state data from the PRIMatE Data Exchange (PRIME-DE) consortium (Milham et al., 2018, n = 25, mean age = 5.4 ± 2.7 years, range 2.4 – 13.1) were used to compute intrinsic functional connectivity patterns in the macaques. These homologous regions derived from the tool-processing ROIs in human adults did not form an intrinsic brain network structure in the macaque brain, as within-tool-domain rsFC was not stronger than between-domain rsFC (*t*_24_ < 0, Figure 2A). In contrast, a face homologous network was observed using the same approach, as within-face-domain rsFC was significantly greater than between-domain rsFC (*t*_24_ = 4.5, *p*_corrected_ < 0.001, *Cohen’s d =* 0.90, Figure 2A). The presence of the face homologous network in macaques was further replicated in a subsample of 9 macaques who were awake during scanning (*t*_8_ = 5.6, *p*_corrected_ = 0.001, *Cohen’s d =* 1.9) and 16 macaques who were anesthetized during imaging acquisition (*t*_15_ = 2.5, *p*_corrected_ = 0.048, *Cohen’s d =* 0.63, Figure S1A).

#### Validation balancing nodal distance and sample size

We first performed a validation analysis based on the left-hemispheric ROIs to address potential confounding effects of nodal distance. The tool processing nodes in the human adults were left-lateralized, whereas the face processing nodes were bilateral. Such differences in hemispheric dominance resulted in greater distances among the tool and face processing nodes compared to the distances among tool processing nodes in all three groups (human adults: *t*_36_ = 2.1, *p* = 0.047; human neonates: *t*_36_ = 2.1, *p* = 0.046; macaques: *t*_36_ = 2.6, *p* = 0.01; the Euclidean distance for each path was generated by averaging from the voxel-to-voxel distances). Such differences in nodal distance might therefore contribute to greater within-tool-domain rsFC compared to the between-domain rsFC. To circumvent this issue, the network effects (i.e., greater within-domain than between-domain rsFC) were further evaluated when the same analyses were carried out with tool and face processing nodes in the left hemisphere only, where the distance of within-tool-domain connections and of between-domain connections were comparable (human adults: *t*_16_ = 0.05, *p* = 0.96; human neonates: *t*_16_ = 0.05, *p* = 0.96; macaques: *t*_16_ = 0.03, *p* = 0.98). The resulting patterns for both the tool and face processing networks remained similar to those in the main analysis. Namely, significantly stronger within-tool-domain rsFC in comparison to between-domain rsFC was still evident in human adults (*t*_99_ = 12.0, *p*_corrected_ < 0.001, *Cohen’s d =* 1.2) and human neonates (for homologous regions, *t*_117_ = 7.6, *p*_corrected_ < 0.001, *Cohen’s d =* 0.70), but not in macaques (for homologous regions, *t*_24_ < 0, Figure 2B). Significantly stronger within-face-domain rsFC compared with between-domain rsFC was observed in human adults (*t*_99_ = 5.6, *p*_corrected_ < 0.001, *Cohen’s d =* 0.56) and macaques (for homologous regions, *t*_24_ = 6.3, *p*_corrected_ < 0.001, *Cohen’s d =* 1.30), but not in human neonates (for homologous regions, *t*_117_ < 0, Figure 2B, see result replications based on subgroups of human neonates and macaques in Figure S1B).

We further considered the potential influences of sample size differences, as the macaque dataset is much smaller (n = 25) than the two human datasets (n ≥ 100) and this was the dataset in which we did not observe the tool homologous network. This concern is assuaged by the fact that we observed the face homologous network in macaques and not in human neonates (i.e., a form of dissociation between the two populations). Nevertheless, a bootstrapping analysis was still conducted to evaluate the network effects among groups with balanced samples (n=25, see Method). As shown in Figure 2C, the distributions of rsFC differences for the tool (homologous) networks in human adults and neonates are greater than 0. As such, the presence of the tool (homologous) network in human adults and neonates is robust across bootstrapping samplings. Note that the bootstrapping results for the face processing network evinced this network’s reliability in human adults (i.e., rsFC differences > 0), but not in human neonates, by corroborating the results reported above using the full sample (Figure 2C).

### Network topology results: Highly similar tool homologous network topology between human adults and neonates, but not between humans and macaques

In addition to the network-level rsFC analyses above, we further characterized and compared every network connection among the tool processing nodes (or their homologues) across the three population groups. Figure 3A visualizes the topological pattern of the tool (homologous) network for each group by showing the path-wise rsFC strengths. The stronger similarity between the human adult and human neonate groups shown in the figure was further confirmed by the topological similarity results on the tool (homologous) processing network (see Figure 1C for the method and Figure 3B for the cross-subject correlation matrix across all subjects). Specifically, the topological patterns of the tool (homologous) networks in human adults and neonates were significantly correlated with large effect sizes (*r* = 0.52 ± 0.48, one-sample *t*_*11799*_ = 119.3, *p*_*corrected*_ < 0.001, *Cohen’s d* = 1.1, Figure 3C). By contrast, the similarities between the macaque group and either human group were, although statistically significant, very low (human adults-macaques: *r* = 0.054 ± 0.49, one-sample *t*_*2499*_ = 5.1, *p*_*corrected*_ < 0.001, *Cohen’s d* = 0.10; human neonates-macaques, *r* = 0.080 ± 0.48, one-sample *t*_*2949*_ = 8.4, *p*_*corrected*_ < 0.001, *Cohen’s d* = 0.16, Figure 3C), and significantly lower than the similarity between the two human groups (human adults-human neonates vs. human adults-macaques: *t*_*14298*_ = 45.0, *p*_*corrected*_ < 0.001, *Cohen’s d* = 0.99; human adults-human neonates vs. human neonates-macaque: *t*_*14748*_ = 46.0, *p*_*corrected*_ < 0.001, *Cohen’s d* = 0.95, Figure 3C).

**Figure 3.**
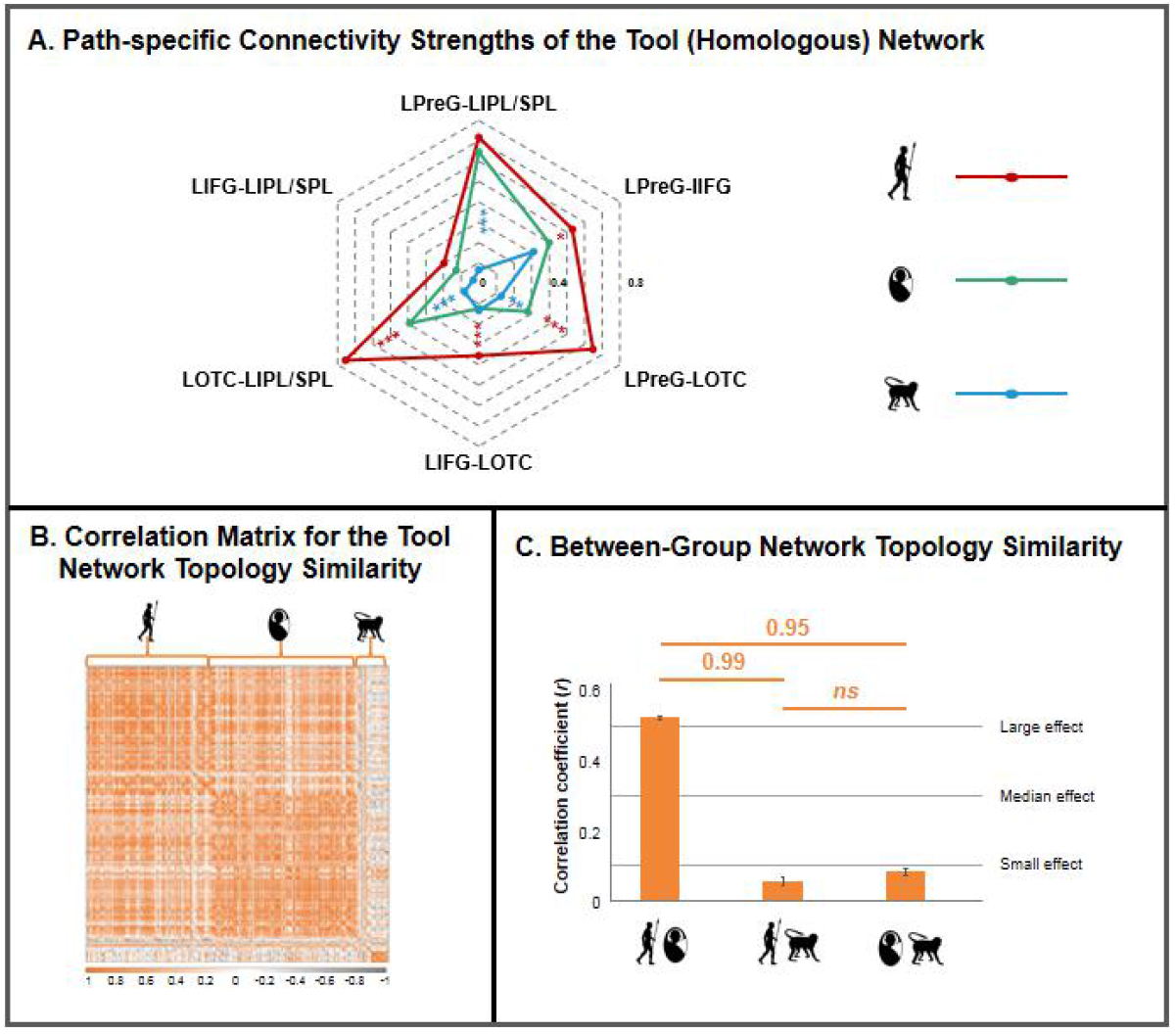
High topology similarities for the tool (homologous) network between human adults and neonates, but not between humans and macaques. A. Path-specific connectivity strengths (Fisher Z scores) of the tool (homologous) network in all three groups. Significant group comparisons between human adults and human neonates are marked in *, whereas significant group differences between human neonates and macaques are marked in *. The left premotor-left inferior/superior parietal path was species-specific, since it was the only path that was comparable between human adults and human neonates, but different between human neonates and macaques. * *p*_*corrected*_ < 0.05, ** *p*_*corrected*_ < 0.01, *** *p*_*corrected*_ < 0.001. B. The correlational matrix for the network topology similarities in all three populations for tool processing networks. C. Bar graphs show tool network topology similarities for participants belonging to different groups. Effect sizes (Cohen’s *d*) are shown for comparisons in which significant differences in between-group pattern similarities were observed (all *p*_corrected_ < 0.001). Error bars indicate standard errors.

For the face homologous network, all between-group correlations for the face (homologous) network were significant with large or medium effect sizes (human adults-neonates: *r* = 0.51 ± 0.26, one-sample *t*_*11799*_ = 234.0, *p*_*corrected*_ < 0.001; human adults-macaques: *r* = 0.47 ± 0.21, one-sample *t*_*2499*_ = 116.7, *p*_*corrected*_ < 0.001; human neonates-macaques, *r* = 0.43 ± 0.25, one-sample *t*_*2949*_ = 100.0, *p*_*corrected*_ < 0.001, Figure S3). The topological patterns for the face (homologous) network were more similar between human adults and neonates than between human adults and macaques (*t*_*14298*_ = 11.1, *p*_*corrected*_ < 0.001, *Cohen’s d* = 0.25), which were in turn were more similar than those between human neonates and macaques (*t*_*5448*_ = 6.6, *p*_*corrected*_ < 0.001, *Cohen’s d* = 0.18, Figure S3). The network topology results mostly remained for both the tool and face processing networks when pre-term and full-term neonates and when awake and anesthetized macaques were analyzed separately (Figure S4).

### Nodal and path results: Strong contributions of premotor connectivity to the formation of the intrinsic tool homologous network in human neonates

Is the formation of the tool (homologous) network in human adults and neonates driven by any particular region(s) or functional connection(s)? This question was addressed using leave-one-node/path-out analyses (Figure 1D). In human adults, when any single node/path was removed, the remaining network still showed stronger within-than between-domain rsFC (all *ts* > 5, all *p*_*corrected*_ < 0.001, Figure S5A), suggesting that the tool processing network was robust in human adults (see the same result patterns derived from the left-hemispheric nodes in Figure S5B). By contrast, in human neonates, when the left premotor node was removed, the remaining tool processing nodes no longer formed an intrinsic network (*t*_*117*_ = 1.2, *p* = 0.24). Removal of any other node or path did not affect the presence of the tool homologous network (i.e., within-tool-domain - between-domain rsFC > 0, all *t*s > 4, all *p*_*corrected*_ < 0.001, Figure 4A). Furthermore, the same analyses were repeated using only the left-hemispheric nodes, to ensure balanced within-tool-domain and between-domain nodal distances. All results were replicated except that the removal of the left inferior/superior parietal node or its connection with the left premotor node made the tool homologous network no longer observable (node removal results: *t*_117_ = 0.24, *p* = 0.81; path removal results: *t*_*117*_ = 1.9, *p* = 0.06, Figure 4B). That is, the connection between left premotor and left inferior/superior parietal nodes is particularly important for the presence of the intrinsic tool homologous network at birth in humans (see replication of the result for nodal and path contributions in subsamples of pre-term and full-term neonates in Figure S6).

**Figure 4.**
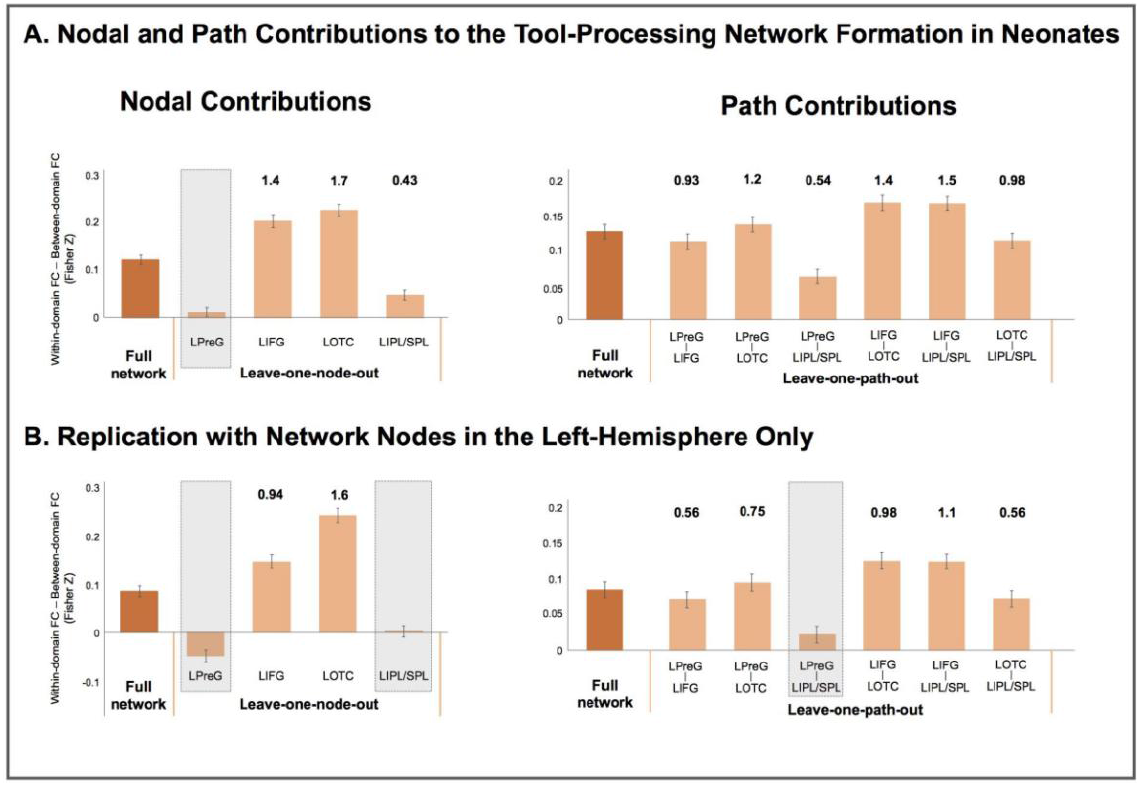
Critical contributions of the premotor region and its connectivity with the parietal region to the formation of the intrinsic tool homologous network in human neonates, as revealed by leave-one-node/path-out analyses. A. Bar graphs illustrate network effects, calculated as within-domain minus between-domain rsFC, for the full tool network and when each of the constituent nodes (left column) or path (right column) is removed. B. Bar graphs exhibit results of the leave-one-node/path out analysis derived from left-hemispheric nodes with balanced within-tool-domain and between-domain path length. Effect sizes (Cohen’s *d*) are shown when the rsFC strengths of the remaining tool network were still significantly higher than those of the between-domain network (all *p*_corrected_ < 0.001). Error bars indicate corresponding standard errors. LOTC: left lateral occipitotemporal cortex; LIPL/SPL: left inferior and superior parietal lobule; LPreG: left premotor gyrus; LIFG: left inferior frontal gyrus; rsFC: resting-state functional connectivity

The path results were further corroborated by the direct group comparisons on the edge-wise rsFC, as visualized in Figure 3A. The left premotor-left inferior/superior parietal connection was the most comparable between human adults and neonates, with smallest *t* value in two-sample samples between human adults and neonates (Table S2) and significantly lower rsFC differences when compared with most of other paths (Table S3). Meanwhile, the LPreG-LIPL/SPL connection also revealed the strongest differences between human neonates and macaques (Figure 3A), with the largest *t* value in two-sample comparisons (Table S2) and significant group x path interaction effects when contrasted with other paths (Table S3). In addition, while the LPreG was significantly connected to the LIPL/SPL in both human adults and neonates (*ts* > 5, *p*_*corrected*_ < 0.001), this connection was not significantly above zero in the macaque brain (*t*_*24*_ = 1.54, *p* = 0.14), further suggesting the species-specific nature of this path.

Note that the non-significant rsFC between LPreG and LIPL/SPL shown in the current macaque dataset seemed to be at odds with previous macaque tracer and fMRI studies (Borra et al., 2017; Howells et al., 2020; Premereur et al., 2015), where the AIP (overlapping with our LIPL/SPL) were found to be connected with the posterior sector of the F5 area (F5p, overlapping with our LPreG, No AIP connections were reliably observed in other two partitions of F5, F5c and F5a). Utilizing the F5 sector masks available in Howells et al. (2020), we observed the positive LIPL/SPL connections in the same F5 sector, F5p (*r* = 0.18, *t*_*24*_ = 6.4, *p* < 0.001), but not in the F5a (*r* = 0.01, *t*_*24*_ = 0.31, *p* = 0.76) or F5c (*r* = 0.01, *t*_*24*_ = 0.23, *p* = 0.82), replicating previous findings. Close comparisons between our registered LPreG ROI and F5 sectors revealed largest overlaps in F5c (34%), compared to F5p (8%), F5a (11%), which likely contributed to the seeming between-study discrepancies noted above. We nonetheless reformed the network analyses in the macaque brain and the between-group comparisons of the LPreG-LIPL/SPL connection with the LPreG ROI replaced by F5p (i.e., the F5 sector with positive rsFC with LIPL/SPL). Again, this path was the most different between human neonates and macaques (Figure S7 and Table S2/S3), and that no tool (homologous) network connectivity structure was observed (within R> between R: *t*_*24*_ < 0). Together, these additional analyses confirmed the species differences in the tool (homologous) network even when different F5 sectors were taken into considerations. The cross-species mapping of the premotor/F5 areas and the different connectivity characteristics of the F5 sectors are theoretically intriguing and were considered further below in Discussion.

## Discussion

To test whether the intrinsic brain connectivity structure supporting tool processing observed in human adults is driven by individual object manipulation experience or is predisposed in humans, we compared their resting-state functional connectivity (rsFC) in this network to homologous networks in human neonates (without manipulation experience) and mature/adolescence macaques (with motor experience with objects), using face (homologous) networks as references. We found that the brain regions that are homologous to those supporting tool processing in human adults were more strongly intrinsically connected with each other than with other nodes (face homologous regions) in the human neonate brain, thereby forming an intrinsic functional network. The homologous regions in macaques did not, however, show a greater within-tool-domain rsFC when compared to the between-domain connectivity. The overall topological patterns among these regions were also highly similar between human adults and human neonates, and much less similar between humans and macaques. The left premotor region, especially its functional connection with the parietal cortex, was particularly important in the formation of the tool homologous network in human neonates.

It should first be acknowledged that the nodes evaluated in human neonate and macaque brains were transformed from regions-of-interest defined in human adult brains using advanced registration methods, including the recently developed cross-species functional alignment approach (Xu et al., 2020) and tools offered by ANTs (Avants et al., 2009). The transformation to neonates’ and other species’ brains is not a trivial task. The cross-species functional alignment approach we adopted here uses a joint-embedding technique that represents the functional organization of human and macaque brains in a high-dimensional common space. This method allows for cortical transformation between species, which had been suggested as the state-of-art transformation approaches (Liu et al., 2021;

Van Essen et al., 2019). Direct comparisons between our transformed ROIs and the manually-drawn regions available in Howells et al., 2020 revealed overlaps between the registered frontal and parietal regions and the key components of the macaque grasping network (Figure S2), confirming the anatomical validity of the current registration approach. Worth noting is that the LIPL/SPL node was mapped onto the AIP and not the IPL region in the macaque brain. AIP has been found to activate when humans and macaques performed hand grasping action and is regarded as an important grasping node, whereas IPL was not typically activated during tool activation viewing in macaques (Borra et al., 2017; Howells et al., 2020; Kastner et al., 2017; Peeters et al., 2009). The registered LIPL/SPL node overlapping with the AIP area in the macaque brain thus reflects the optimal performance of the current cross-species functional alignment approach in maximizing the functional similarity between the source and target brain spaces. It is, therefore, unlikely that the absence of the tool homologous network in macaques is due to transformation problems. Next, is it possible that within-tool-domain rsFC was stronger than between-domain rsFC in human neonates due to registration errors in neighboring networks supporting other types of cognition, and not due to anything related to tools? Resting-state functional connectivity studies on early development have identified large-scale brain networks in infancy using parcellation approaches, and the networks reported in these studies do not overlap with the tool homologous network we observed here (Doria et al., 2010; Fransson et al., 2007; Gao et al., 2015, except for Fitzgibbon et al., 2020, in which one of the 16 sub-networks is visually similar). Future task-based fMRI and/or neurophysiological studies with infant and macaque populations are warranted to further elucidate nodal function. The rsFC findings in the present study are well-suited to guide such future studies, given the tight coupling between connection and function (connection determines function, Mars et al., 2018; Passingham et al., 2002).

Our observation that adolescent/mature macaques do not show strong intrinsic connectivity among regions homologous to tool processing nodes in human adults suggests that the tool processing network may be (at least partly) specific to humans. This network is not (fully) driven by simple associations across sensory-motor experiences, as these mature and adolescent macaques have ample experience with objects. This observation also provides a neural basis for germane cognitive differences between the two species. Namely, while macaques are capable of grasping and using simple tools based on surface property associations, they lack an understanding of the causal mechanic mechanisms related to tools and are incapable of generalizing functions of objects (Vaesen, 2012). Therefore, intrinsic functional connectivity among tool homologous regions, at least in part, seems to support the casual (mechanic) reasoning exhibited by humans, but not macaques. This interpretation gains further support from the most significant differences in the frontal to parietal connection observed between humans and macaques. Such findings are consistent with the cross-species differences in the (putative) functional characteristics of these two regions with respect to tool-related activation. The IPL region has been shown to exhibit preferences for viewing tool actions (as compared to hand grasping actions) only in humans, whereas no such functional preferences could be reliably observed in the parietal cortex of macaques (Peeters et al., 2009). Instead, its FC-based homologous region (AIP) were typically activated during the hand grasping movement in macaques (Kastner et al., 2017). AIP has further been shown to connect with the posterior partition of the F5 (F5p), and together they formed a neural network thought to support hand grasping abilities of macaques (Howells et al., 2020; Premereur et al., 2015). Moreover, even though the left premotor area in the human brain could be generally mapped to the F5 region of the macaque brain, it overlapped the most with the convexity sector, F5c. Limited findings seem to suggest different effector bias between different F5 sectors: the F5c, and F5a to a lesser extent, respond almost exclusively to hand-related action, whereas the F5p is also involved in the mouth movement with selective neurons discharging during monkey vocalization (Borra et al., 2017). These results together suggest different functional relevance for the homologous regions between two species, reflecting the intrinsic differences in the tool-processing network in humans and the grasping network in macaques.

For human neonates, the functionality and connectivity patterns of regions that become tool-sensitive later in life are even more poorly understood. To date, most studies investigating rsFC in infants have focused on primary (e.g., sensorimotor and visual) functional networks and higher-order functional networks underlying domain-general cognition (Gilmore et al., 2018), and have reported protracted maturation of the default-mode and attention networks such that they do not reach adult-like topology until age two (Fiske and Holmboe, 2019; Gao et al., 2015). Our observations that the tool homologous network is already established in human neonates, and that the intrinsic functional connectivity pattern of this network is more similar between human adults and human neonates than between human adults and macaques, who have plenty of object perception/grasping experience, indicate that the network supporting tool processing is innate and in place early in human development. Given that neonates have not yet developed any voluntary motor skills, the presence of the tool network at this developmental stage indicates that its formation does not depend upon individual object (tool) manipulation experience. Furthermore, the functional connection between the premotor and parietal clusters is particularly important in driving the formation of this network in human neonates. Taken together, this network, especially the premotor-parietal connection, offers a brain basis for the development of causal (mechanic) understanding of using objects as tools in humans (and not macaques). This speculation is in line with the contemporaneous emergence of flexible tool use behaviors and perceptual and motor skills at early developmental stages (Kastner et al., 2017), and with lesion studies in human adults, which showed significant associations between damage to parietal-premotor white matter connections and difficulty understanding and using tools (Bi et al., 2015). Further studies are invited to further elucidate the white matter structural connectivity in human neonates and nonhuman primates, to determine whether and to what extent development of tool use and training with tools may modulate the tool processing network, and to identify the function of various components of the neonate tool (homologous) network.

Finally, our results for the face homologous network in human neonates are worth discussing. While the overall connectivity topology among the face (homologous) nodes was significantly similar between human adults and neonates, more so than that between human and macaque species, we did not observe a robust intrinsic face homologous network based on within-face-domain versus between-domain comparisons. That is, the face homologous regions were not robustly segregated from the tool homologous nodes in neonates. Previous studies have reported face-relevant profiles similar to adults within the visual cortex of human neonates/infants: similar to adults, intrinsic functional connections among the homologues of occipital face areas, fusiform face areas, and foveal V1 areas were observed in neonates (Kamps et al., 2020); also similar to adults, face-preferring topography was observed in the ventral visual cortex during face-viewing in infants 4-6 months of age (Deen et al., 2017). Together, these results suggest that the intrinsic functional topology potentially supporting face processing is at least partly present at birth, and visual experience may be needed in developing a more functionally segregated large-scale face processing network beyond the visual cortex (see also Arcaro et al., 2017).

To conclude, for the brain network supporting tool processing in human adults, the intrinsic functional connectivity network structure is absent among the homologous regions in our evolutionary cousins, macaque monkeys, but is present in humans at birth before they have had individual experiences interacting with objects. The functional connection between premotor and parietal nodes, in particular, is important for the formation of the tool homologous network connectivity in human neonates and thereby constitutes a strong candidate for the neural basis of complex tool use specific to humans. These results contribute empirical evidence to the broad issue of which neural aspects are human specific, and highlight the need for further research to understand the neural and computational underpinnings of tool use.

## Materials and Methods

### Participants

#### Human adults

Resting-state images of human adults were obtained from the WU-Minn Human Connectome Project (HCP) carried out at Washington University in St. Louis (Van Essen et al., 2013, https://www.humanconnectome.org/study/hcp-young-adult). For the current study, 100 individuals (55 females, 28.3 ± 3.4 years old), coming from different families (i.e., not related), were randomly selected from the 1200 Subjects Data Release. The fMRI data of all selected participants met the following inclusion criteria: 1) had less than 10% of volumes with framewise displacement ≥ 0.3 mm (see details in **Image preprocessing**) and 2) exhibited good coverage of the functional ROIs selected (see details in **Functional ROI selection**). This project was reviewed and approved by the Institutional Ethics Committee of Washington University in St. Louis, Missouri. All participants signed written informed consent.

#### Human neonates

Imaging data of human neonates were obtained from the Developing Human Connectome Project (dHCP) conducted at the Newborn Imaging Centre at Evelina London Children’s Hospital, London, UK (Makropoulos et al., 2018, https://www.developingconnectome.org). 118 neonates (57 females, birth age = 39.7 ± 1.9 weeks; scan age = 40.9 ± 2.1 weeks, birth weight = 3.1 ± 0.66 kg) were selected from the two data releases available at the time of data analysis for the present study based on the following inclusion criteria: 1) images were acquired within the first month (i.e., ≤ 4 weeks) after birth; 2) structural images showed no clinical concerns when evaluated by a perinatal neuroradiologist (i.e., radiology score ≤ 3); 3) ≤ 10% of scans contained excessive head movement, defined as ≥ 0.3 mm framewise displacement; 4) there was good coverage of the functional ROIs selected. Among them, 12 neonates were born pre-term (birth age range: 31.7-36.9 weeks), while the remaining 106 participants were born full-term. The dHCP was approved by the UK health Research Authority. Informed parental consent was obtained for imaging acquisition and data release.

#### Adolescent and adult macaques (macaca mulatta)

Macaque imaging data were obtained from the PRIMatE Data Exchange (PRIME-DE) consortium (Milham et al., 2018, http://fcon_1000.projects.nitrc.org/indi/indiPRIME.html). Two cohorts of macaques were included in the present study. One of them was *Newcastle data*, where macaques were *awake* during resting-state image acquisition. Resting-state fMRI images were available for 10 macaques (Baumann et al., 2015, 2011; Poirier et al., 2017; Rinne et al., 2017; Schönwiesner et al., 2015; Slater et al., 2016; Wilson et al., 2015), but data from one macaque was removed due to poor coverage of the functional ROIs, resulting in 9 macaques (2 females, age = 8.4 ± 2.5 years, weight = 11.6 ± 3.6 kg) whose data were entered in the final analysis. The other one was *Oxford data*, where macaques were *anesthetized* during imaging data collection. The original dataset consisted of 20 rhesus macaque monkeys (Noonan et al., 2014). However, data of four macaques were excluded due to poor normalizations (n = 1) or insufficient coverage of functional ROIs (n = 3), resulting in a final set of 16 macaques (all males, age = 3.7 ± 0.69 years, weight = 5.9 ± 1.4 kg). Together, there were 25 macaques (n = 25, mean age = 5.4 ± 2.7 years, range 2.4 – 13.1) whose fMRI images were included in the current study (see supplementary Table S1 for the full list of subject IDs).

### Image acquisition

#### Human adults

Images were collected using a 3T Siemens Skyra magnetic resonance scanner with a 32-channel head coil (Van Essen et al., 2013). Resting-state images were collected while participants fixated (eyes open) on a bright cross-hair projected on a dark background (and presented in a darkened room). A gradient-echo echo planar imaging (GE-EPI) sequence was applied with the following parameters: repetition time (TR) = 720 ms, echo time (TE) = 33.1 ms, flip angle (FA) = 52°, bandwidth = 2290 Hz/pixel, field of view (FOV) = 208 × 180 mm_^2^_, matrix = 104 × 90, voxel size = 2 × 2 × 2 mm^3^, multi-band accelerated factor = 8, slices = 72, and total scan time of 1200 frames = 14 min and 33s (Smith et al., 2013). Two sessions (i.e. REST1 and REST2) were collected on two consecutive days, each including two runs with both phase encoding directions (i.e. left-to-right and right-to-left). All four runs were used in the present study. High-resolution T1-weighted images were also acquired for every participant using a magnetized rapid gradient-echo imaging (MPRAGE) sequence with TR = 2400 ms, TE = 2.14 ms, reversal time (TI) = 1000 ms, FA = 8°, FOV = 224 × 224 mm^2^, voxel size = 0.7 mm isotropic, and total scan time = 7 min and 40 s.

#### Human neonates

Images were collected using a 3T Philips Achieva with a dedicated neonatal imaging system, including a neonatal 32-channel phased array head coil (Hughes et al., 2017). All neonates were scanned during nature sleep without sedation. A multiband EPI sequence was utilized with TR = 392 ms, TE = 38ms, FA = 34°, voxel size = 2.15 × 2.15 × 2.15 mm^3^, MB factor = 9x, and total scan time = 2300 volume (15min and 3.5s). T2-weighted (TR = 12 s; TE = 156 ms; SENSE factor: axial = 2.11, sagittal = 2.58) and inversion recovery T1-weighted (TR = 4795 ms; TI = 1740 ms; TE = 8.7 ms; SENSE factor: axial = 2.26, sagittal = 2.66) multi-slice fast spin-echo images were also collected for each neonate (in-plane resolution = 0.8 × 0.8 mm^2^, 1.6 mm slices overlapped by 0.8 mm, see details in Fitzgibbon et al., 2020).

#### Macaque monkeys (macaca mulatta)

The *Newcastle data* were collected on a Vertical Bruker 4.7T primate dedicated scanner with a single channel or a 4-8 channel parallel imaging coil. All monkeys included in the current study were awake during resting-state imaging acquisition. Two resting-state sessions were collected for each monkey with 1.2 × 1.2 × 1.2 mm^3^ resolution, TR = 2600 ms, TE = 17 ms, 10.8-min (250 volumes) per scan. T1-weighted images were also acquired using a MDEFT sequence with the following parameters: TR = 750 ms, TE = 6ms, inversion delay = 700 ms, FOV = 12.8 × 9.6 cm^2^ on a grid of 256 × 192 voxels, voxel size = 0.5 × 0.5 × 2 mm, number of slices = 22. Additionally, no contrast agent was used during scanning.

The *Oxford data* were collected on a 3T scanner with a 4-channel coil when macaques were under anesthesia. Again, no contrast agent was used. The acquisition parameters for the resting-state images were 2 × 2 × 2 mm^3^ resolution, TR = 2s, TE =19 ms, FP = 90°, and total scan time = 53.3 min (1600 volumes). T1-weighted images for each monkey were also acquired using a MPARGE sequence with the following parameters: TR = 2500 ms, TE = 4.01ms, TI = 1100 ms, FP = 8°, voxel size = 0.5 × 0.5 × 0.5 mm.

### Image preprocessing

#### Human adults

We used the HCP’s minimally preprocessed resting-state data (Glasser et al., 2013), which were distortion and motion corrected and registered to MNI templates via structural images using non-linear transformations. These images were further denoised using independent component analysis (ICA) with the FIX tool (Salimi-Khorshidi et al., 2014) to effectively identify and remove the components of spatiotemporal signals caused by non-neuronal or structural noise, including head movement (Smith et al., 2013). Moreover, volumes with ≥ 0.3 mm framewise displacement (FD, Power et al., 2012) were identified as outlier scans with excessive motion. All human adults included in the current analyses had no more than 10% outliers (2.7% ± 0.025). Preprocessing procedures subsequently performed using the DPABI toolbox (Yan et al., 2016) included: 1) linear detrending to minimize the effects of low-frequency drift; 2) regression of nuisance variables, including the mean white matter (WM) and the cerebrospinal fluid (CSF) signals, continuous head movement (Friston-24 parameters, Friston et al., 1996) and outlier scans, to further reduce non-neuronal contributions, 3) temporal band-pass (0.01-0.1 Hz) filtering to decrease non-neurophysiological noise, and 4) spatial smoothing (Gaussian filter, FWHM = 6 mm).

#### Human neonates

Similar to the HCP dataset, the resting-state functional images of the neonates first underwent a minimally preprocessed pipeline developed by the dHCP team specifically for this age range (Fitzgibbon et al., 2020). This pipeline included motion and distortion correction, registration of the functional images with corresponding T2-weighted images, as well as ICA-FIX denoising. Deformational matrices for aligning individual structural images in native space to a 40-week T2 template were also generated, and subsequently applied to the minimally preprocessed functional images to normalize them to the 40-week standard space. Images with excessive motion, defined as ≥ 0.3 mm FD, were identified and all neonates included in the current analyses had few than 10% of outliers (5.1% ± 0.028). Similar to the human adults, the images of the neonates were subsequently preprocessed using the DPABI toolbox for linear detrending, removal of nuisance effects (mean WM and CSF time series, Friston-24 parameters and outlier scans), temporal band-pass filtering (0.01-0.1Hz), and spatial smoothing (Gaussian filter, FWHM = 6 mm).

#### Macaque monkeys

Images of both awake (*Newcastle data*) and anesthetized (*Oxford data*) monkeys were fully preprocessed using the DPABI toolbox with the following steps: 1) discarding the first five time points for signal equilibrium and adaptation to the scanning noise, 2) correcting for head movement, 3) removing the signal trend linearly, 4) identifying outliers defined as ≥ 0.3 mm FD, 5) regressing out the nuisance variables, including the mean WM and CSF time series, continuous head movement and outlier scans, 6) normalizing to the 112RM-SL template (the volume-based atlas (McLaren et al., 2010, 2009) using unified segmentation on T1-weighted images, 7) band-pass (0.01–0.1 Hz) filtering, and 8) spatial smoothing with a 3 mm full-width half-maximum Gaussian kernel. The anesthetized macaques showed minimal head movement during scanning with no outlier images. However, the awake macaques showed a variable number of outlier images (3%-29%, mean = 13% ± 0.11). Due to the small sample of awake macaques, their images were still included in the current study. Nevertheless, main results obtained based on the whole macaque group were replicated in awake and anesthetized macaques separately, ensuring the reliability of the current findings.

### Functional ROI selection and cross-population registration (Figure 1A)

Nodes (regions-of-interest; ROIs) of the neural networks underlying tool (and face as a control domain) processing in human adults were objectively generated from meta-analyses based on the Neurosynth database incorporating 14,371 fMRI studies in total (https://neurosynth.org, version 0.7 released July, 2018, Yarkoni et al., 2011). Association maps based on the terms “tools” and “face” were generated respectively using the default threshold at FDR corrected, p < 0.01 with *k* = 50 voxels. The tool processing network, derived from 115 studies, contained three regions in the left hemisphere: left lateral occipitotemporal cortex (LOTC), left inferior frontal gyrus (LIFG), and left inferior and superior parietal lobule (LIPL/SPL). Given the frequently reported involvement of the left premotor area (LPreG) in tool-relevant tasks (Brandi et al., 2014; Lewis, 2006), including simply viewing (Chao and Martin, 2000), an additional ROI located in the LPreG was further obtained using a lenient threshold at z = 3.09. The face processing network was derived from 896 studies and initially revealed five cerebral ROIs, including LSTG, RIFG, and three large clusters in the left (*k* = 3229 voxels) and right (*k* = 4728 voxels) ventral visual pathways, as well as the RATL, which extended into the subcortical areas (*k* = 1156 voxels). For the two clusters in the ventral visual pathways, more stringent thresholds were applied to identify functionally distinctive ROIs, resulting in RFFG, ROFA and RSTG on the right (z = 5) and LFFA and LOFA on the left (z = 8). The stricter threshold (z = 5) also helped to confine the RATL to the cerebral cortex. Overall, we identified a left-hemispheric tool network and a bilateral face network (Figure 1A) that contained key regions commonly reported in previous meta-analyses and review papers (Lewis, 2006; Wang et al., 2020).

For neonates, these tool and face processing ROIs identified in human adults in MNI152 space were then transformed onto 40-week templates available on the developmental human connectome project (dHCP) website (https://gin.g-node.org/BioMedIA/dhcp-volumetric-atlas-groupwise), using ANTs (Figure 1A). Hereafter, the transformed ROIs were referred to as tool (or face) homologous nodes or regions, emphasizing that they are brain regions homologous to those showing tool processing sensitivity in human adults, and not directly functionally defined in human neonates (and macaques).

For macaques, registration of the ROIs was achieved using the landmark-based functional connectivity approach recently developed by (Xu et al., 2020). Specifically, volumetric ROIs identified in human adults for tool and face processing were first mapped onto a standard surface (i.e., 32k_fs_LR) using the registration fusion approach (Wu et al., 2018). They were then transferred from the human surface to the macaque space (Yerkes 19 atlases, Donahue et al., 2016) using a joint-embedding technique. This approach represents the functional organization of macaques and humans in a high-dimensional common space which enables establishing the cortical transformation between these two species (Xu et al., 2020). The transformed ROIs, now in macaque surface space, were then converted into volumetric space using the HCP workbench command (label-to-volume-mapping, ribbon constrained mapping algorithm) and registered to the volume-based 112RM-SL template (DPABI defaults) using ANTs. The registration results were visually inspected and validated by computing overlaps to the hand-drawn ROIs in the macaque brain available in a previous study (see results).

Finally, to ensure sufficient coverage of the selected ROIs for the FC analyses, a binary brain mask was generated based on the preprocessed functional images of each participant using the DPABI automask function. The overlap between the analysis mask and each ROI was calculated. All the included human adults and neonates showed optimal coverage of each ROI with a minimum of 50% overlap (adults: 96% ± 0.06, neonates: 96% ± 0.09). A lenient threshold (30%) was applied to the macaque groups to maximize the sample sizes of the awake (95% ± 0.12) and anesthetized (96% ± 0.13) macaques. Nevertheless, replication results were obtained based on the data of 17 macaques who met the 50% coverage threshold comparable to the human participants, ensuring that the observed effects cannot be attributed to poorer coverage of the functional ROIs in macaques’ images (Figure S8).

### Resting-state functional connectivity (rsFC) analyses

For each individual from all the three groups (i.e. human adults, human neonates, and macaques), the node-based timecourse was calculated by averaging across all voxels included in each ROI. Pearson correlations were then performed for each ROI pair and resulting correlation coefficients were normalized to z-scores using Fisher’s z transformation. This procedure generated an rsFC matrix for each subject. Network analyses included three major steps. First, the intrinsic tool and face processing networks were evaluated in each group by comparing the rsFC between nodes belonging to the same domain with that between nodes from different domains using paired *t*-tests (Figure 1B). An intrinsic network was established if the within-domain rsFC was significantly greater than the between-domain rsFC. Two validation analyses were subsequently performed to ensure the observed network effects were independent of confounding variables of nodal distance and sample size. Specifically, the tool processing nodes in the human adults were left-lateralized, whereas the face processing nodes were bilateral, resulting in greater distances in the connectivity between the tool and face processing nodes (i.e. between-domain path) compared to the distances among tool processing nodes in all three groups (all *ps* < 0.05). Such differences in nodal distance might contribute to greater within-tool-domain rsFC compared to the between-domain rsFC, if observed. To circumvent this issue, the network effects (i.e., greater within-domain than between-domain rsFC) were further evaluated when the same analyses were carried out with tool and face processing nodes in the left hemisphere only, where the distance of within-tool-domain connections and of between-domain connections were comparable (all *ps* > 0.9). Next, a bootstrapping analysis was performed to deal with the sample size differences, as the macaque dataset was much smaller (n = 25) than the two human datasets (n ≥ 100). To do this, the subsamples of human adults and neonates with the subject sizes equal to that of macaque sample were created by randomly sampling 25 data from the HCP and dHCP datasets, respectively. These resting-state images were submitted to the same network analyses as described in the main analysis for comparisons of within-domain and between-domain rsFC. This process was repeated 10,000 times, producing group-specific distributions of within-domain minus between-domain rsFC. The network effect obtained based on the macaque dataset was then compared against these distributions of the network effects derived from the subsamples of human adults and neonates with equal sample sizes.

The second analysis focused on the network topology similarity between different groups. Pearson correlations were conducted on the rsFC values across paths within the tool (or face) processing network for each subject pair across all three groups, which were converted into Fisher Z scores (Figure 1C). One-sample *t*-tests were applied to evaluate whether each of the between-group pattern similarities were significantly greater than 0. Two-sample *t*-tests were subsequently conducted to compare between-group similarities among different group pairs.

In the final analysis, the contribution of each node and each path to the intrinsic tool network observed in human adults and neonates was investigated. A leave-one-node/path-out approach was applied, where the comparisons of within- and between-domain rsFC were re-evaluated when one node or path was removed at a time (Figure 1D). Correction for multiple comparisons was performed for every analysis. The Cohen’s *d* effect size was additionally computed for the *t*-test results for clearer interpretation. For correlation measures, after statistical analyses, z-scores were transformed back to *r* values for illustration purposes.

## Acknowledgments

This work was supported by the National Natural Science Foundation of China (32100867 to X.Y., 31925020, 31671128 to Y.B., 32071050 to X.Y.W), Changjiang Scholar Professorship Award (T2016031 to Y.B.), the National Program for Special Support of Top-Notch Young Professionals (to Y.B.). The human adult data were provided by the Human Connectome Project, WU-Minn Consortium (Principal Investigators: David Van Essen and Kamil Ugurbil; 1U54MH091657), which is funded by the 16 NIH Institutes and Centers that support the NIH Blueprint for Neuroscience Research; and by the McDonnell Center for Systems Neuroscience at Washington University. The neonate data were provided by the developing Human Connectome Project, KCL-Imperial-Oxford Consortium, which is funded by the European Research Council under the European Union Seventh Framework Programme (FP/2007-2013) / ERC Grant Agreement no. [319456]. The macaque data were provided by investigative teams from Oxford (Principal Investigators: Jerome Sallet, Rogier B. Mars, Matthew F.S. Rushworth) and funded by the Wellcome Trust, Royal Society, Medical Research Council UK and the Biotechnology Biological Sciences Research Council UK, and the teams from Newcastle (Principal Investigators: Jennifer Nacef, Christopher I. Petkov, Fabien Balezeau & Timothy D. Griffiths, Colline Poirier & Alexander Thiele, Michael Ortiz & Michael Schmid, David Hunter), which are funded by the Wellcome Trust, National Center for 3Rs, UK Biotechnology Biological Sciences Research Council and National Institutes of Health. We would like to thank the investigative teams providing these publicly available datasets and the funding agencies that make these datasets available. We would also like to acknowledge the assistance of Yaya Jiang, Yumeng Xin and Xinyu Liang for constructive suggestions in data analysis, and Xinyi Tang for helpful comments on earlier drafts of the manuscript.

## Author contributions

Haojie Wen: Conceptualization, Methodology, Formal analysis, Investigation, Writing, Writing-original draft, Visualization. Ting Xu: Methodology, Writing-Review and Editing. Xiaoying Wang: Conceptualization, Methodology, Writing-Review and Editing, Funding acquisition. Xi Yu: Conceptualization, Methodology, Formal analysis, Writing-original draft, Review and Editing, Visualization, Supervision, Funding acquisition. Yanchao Bi: Conceptualization, Methodology, Writing-original draft, Review and Editing, Supervision, Funding acquisition.

## Data and code availability statement

The human adult and neonate datasets are available at https://www.humanconnectome.org/ and https://www.developingconnectome.org, respectively. The macaque data are available at http://fcon_1000.projects.nitrc.org/indi/indiPRIME.html. For the macaque data, subject IDs included in the current study are listed in Supplementary Table S1, and the computed network results are available at https://github.com/xiyu-bnu/neonate_tool_network. Due to HCP and dHCP privacy policies, the preprocessed resting-state images of human adults and neonates (with their IDs) can only be shared upon request with qualified investigators who agree to the Restricted Data Use Terms of these two datasets.

## Supplementary Information

**Table S1.** A full list of the macaque subject IDs included in the current analysis.

| ID in the current project | Dataset | Original subject ID |
| --- | --- | --- |
| Macaque 001 | Newcastle | 32097 |
| Macaque 002 | Newcastle | 32100 |
| Macaque 003 | Newcastle | 32104 |
| Macaque 004 | Newcastle | 32105 |
| Macaque 005 | Newcastle | 32106 |
| Macaque 006 | Newcastle | 32107 |
| Macaque 007 | Newcastle | 32108 |
| Macaque 008 | Newcastle | 32109 |
| Macaque 009 | Newcastle | 32110 |
| Macaque 010 | Oxford | 32163 |
| Macaque 011 | Oxford | 32165 |
| Macaque 012 | Oxford | 32166 |
| Macaque 013 | Oxford | 32167 |
| Macaque 014 | Oxford | 32169 |
| Macaque 015 | Oxford | 32171 |
| Macaque 016 | Oxford | 32172 |
| Macaque 017 | Oxford | 32173 |
| Macaque 018 | Oxford | 32174 |
| Macaque 019 | Oxford | 32175 |
| Macaque 020 | Oxford | 32176 |
| Macaque 021 | Oxford | 32177 |
| Macaque 022 | Oxford | 32178 |
| Macaque 023 | Oxford | 32179 |
| Macaque 024 | Oxford | 32180 |
| Macaque 025 | Oxford | 32181 |

**Table S2.** Statistical results of cross-population comparisons for path-specific functional connectivity.

|  | LPreG-LIFG |  |  | LPreG-LOTG |  |  | LPreG-LIPL/SPL |  |  | LIFG-LOTG |  |  | LIFG-LIPL/SPL |  |  | LOTG-LIPL/SPL |  |  | df |
| --- | --- | --- | --- | --- | --- | --- | --- | --- | --- | --- | --- | --- | --- | --- | --- | --- | --- | --- | --- |
|  | Dif | t | p | Dif | t | p | Dif | t | p | Dif | t | p | Dif | t | p | Dif | t | p |  |
| **Human adults VS. Neonates** | 0.13 | 3.8 | * | 0.37 | 12.5 | *** | 0.07 | 1.9 | (0.054) | 0.23 | 8.7 | *** | 0.07 | 2.4 | (0.02) | 0.36 | 11.7 | *** | 216 |
| **Human neonates VS. Macaques** | 0.09 | 1.6 | (0.11) | 0.15 | 3.2 | ** | 0.58 | 9.9 | *** | -0.01 | -0.26 | (0.80) | 0.10 | 2.2 | (0.03) | 0.31 | 6.2 | *** | 141 |
| **Human neonates VS. Macaques**<br>**(replacing LPreG by F5p)** | 0.17 | 3.3 | ** | 0.15 | 3.2 | * | 0.47 | 8.3 | *** | -0.01 | -0.26 | (0.80) | 0.10 | 2.2 | (0.03) | 0.31 | 6.2 | *** | 141 |

Note. \* indicates $p_{corrected} < 0.05$ after Bonferroni correction ( $n = 6$ ). For non-significant results, the uncorrected $p$ is presented in parentheses.
\* $p_{corrected} < 0.05$ , \*\* $p_{corrected} < 0.01$ , \*\*\* $p_{corrected} < 0.001$ .

**Table S3.** Interaction effects of group (human neonates vs. other groups) × path (LPreG-LIPL/SPL vs. other paths)

|  | LPreG-LIFG |  | LPreG-LOTG |  | LIFG-LOTG |  | LIFG-LIPL/SPL |  | LOTG-LIPL/SPL |  | df |
| --- | --- | --- | --- | --- | --- | --- | --- | --- | --- | --- | --- |
|  | *F* | *p* | *F* | *p* | *F* | *p* | *F* | *p* | *F* | *p* |  |
| **Human adults VS. Neonates** | 1.1 | (0.29) | 76.7 | * | 14.4 | * | 0.01 | (0.92) | 55.7 | * | 1, 216 |
| **Human neonates VS. Macaques** | 35.8 | * | 73.7 | * | 105.3 | * | 80.0 | * | 21.6 | * | 1, 141 |
| **Human neonates VS. Macaques**<br>**(replacing LPreG by F5p)** | 13.9 | * | 47.0 | * | 69.6 | * | 45.5 | * | 8.07 | * | 1, 141 |

Note. \* indicates $p_{corrected} < 0.05$ after Bonferroni correction ( $n = 5$ ). For non-significant results, the uncorrected $p$ is presented in parentheses.
LOTG: left lateral occipitotemporal cortex; LIPL/SPL: left inferior and superior parietal lobule; LPreG: left premotor gyrus; LIFG: left inferior frontal gyrus

**Figure S1.**
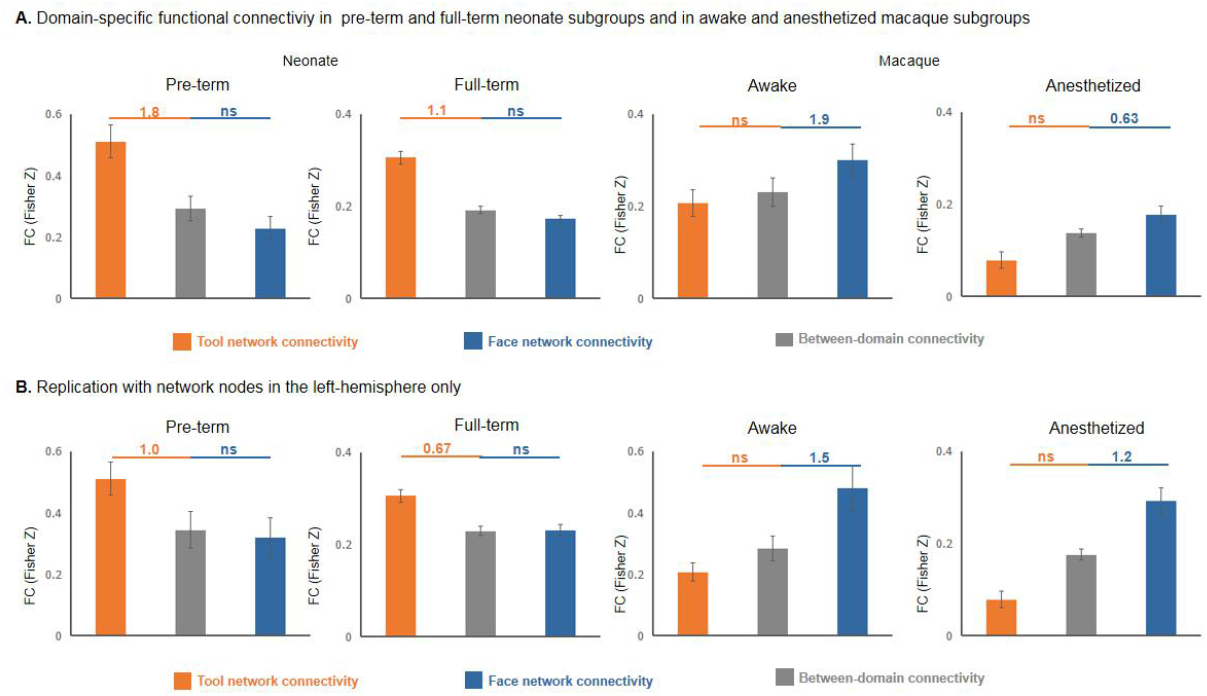
Resting-state functional connectivity (rsFC) results when the pre-term neonates, full-term neonates, awake macaques, and anesthetized macaques were analyzed separately. A. Bar graphs illustrate the rsFC values for within- and between-domain functional connectivity in the pre-term neonate, full-term neonate, awake macaque, and anesthetized subgroups. B. Bar graphs show replication results for the network analysis restricted to the left-hemispheric nodes with balanced within-tool-domain and between-domain nodal distance. Within-domain rsFC values were greater than between-domain rsFC values in pre-term (*t*_*11*_ = 3.6, *p*_*corrected*_ = 0.008, *cohen’s d* = 1.0) and full-term neonates (*t*_*105*_ = 6.9, *p*_*corrected*_ < 0.001, *cohen’s d* = 0.67) for the tool processing network, but not for the face processing network (pre-term: *t*_*11*_ < 0; full-term: *t*_*105*_ = 0.20). Within-domain FC values were greater than between-domain FC values in awake (*t*_*8*_ = 4.6, *p*_*corrected*_ = 0.004, *cohen’s d* = 1.5) and anesthetized macaques (*t*_*15*_ = 4.7, *p*_*corrected*_ < 0.001, *cohen’s d* = 1.2) for the face processing network, but not for the tool processing network (both *t* < 0). Effect sizes (Cohen’ s *d*) are shown for comparisons with significant differences (all *p*_*corrected*_ < 0.001, Bonferroni corrected (n = 2)). Error bars indicate standard errors.

**Figure S2.**
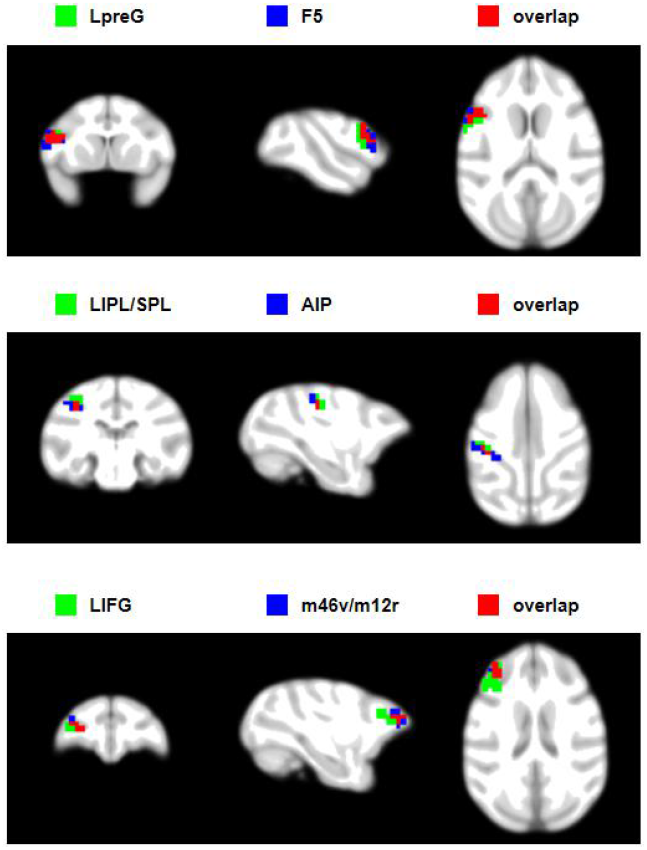
Overlaps (red) between the tool homologous nodes (green) in the macaque brain and the manually-drawn lateral grasping network (blue) available in Howells et al. (2020). The left premotor gyrus (LPreG) and F5 are presented in the top panel, the left inferior and superior parietal lobule (LIPL/SPL) and AIP in the middle panel, and the left inferior frontal gyrus and m46v/m12r in the bottom panel.

**Figure S3.**
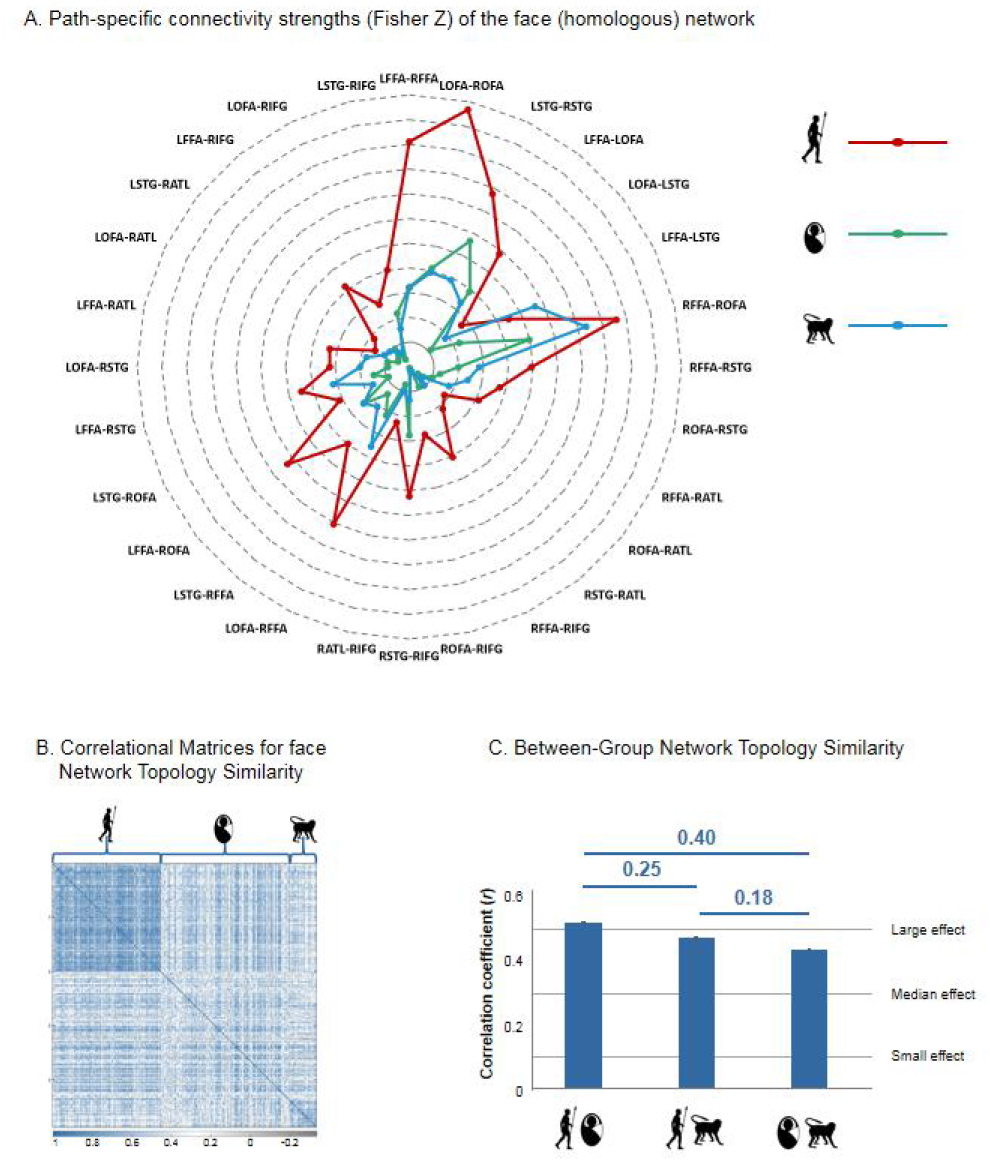
The topological pattern for the face (homologous) network. A. Path-specific connectivity strengths (Fisher-Z scores) of the face (homologous) network in all three groups. All group comparisons between human adults and human neonates were significant (all *t* > 5, *p*_*corrected*_ < 0.001, Bonferroni corrected (n = 28)). Most of the paths were comparable between human neonates and monkeys, except LFFA-LSTG (*t* = -7.8, *p*_*corrected*_ < 0.001), LOFA-RFFA (*t* = -3.4, *p*_*corrected*_ = 0.03), RFFA-ROFA (*t* = -4.8, *p*_*corrected*_ < 0.001), LFFA-RSTG (*t* = -4.8, *p*_*corrected*_ < 0.001), LOFA-RSTG (*t* = -3.3, *p*_*corrected*_ = 0.03), LSTG-RSTG (*t* = 3.4, *p*_*corrected*_ =0.001). B. The correlational matrix for the face (homologous) network topology similarity among all three population groups. C. Bar graphs show face network topology similarities for participants belonging to different groups. Effect sizes (Cohen’s *d*) are shown for comparisons in which significant differences in between-group pattern similarities was observed (all *p*_corrected_ < 0.001). Error bars indicate standard errors.

**Figure S4.**
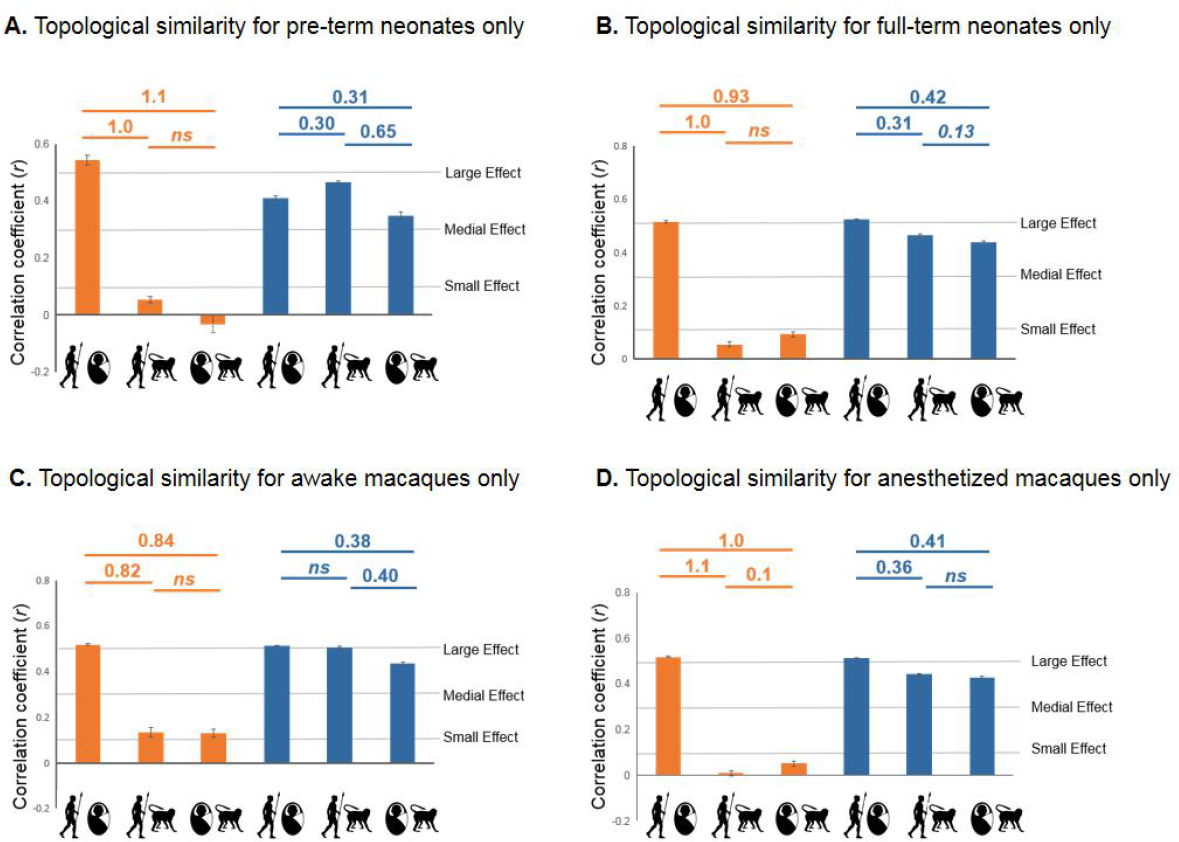
Network topology similarity results in subgroups of pre-term (A) and full-term (B) neonates, as well as awake (C) and anesthetized (D) macaques. The results of subgroups remained unchanged in each analysis. Bar graphs illustrate the network topological pattern similarities for participants belonging to different groups. Effect sizes (Cohen’s *d*) are shown for comparisons existing significant differences in the between-group pattern similarities (all *p*_corrected_ < 0.05, Bonferroni corrected (n = 6)). Error bars indicate standard errors.

**Figure S5.**
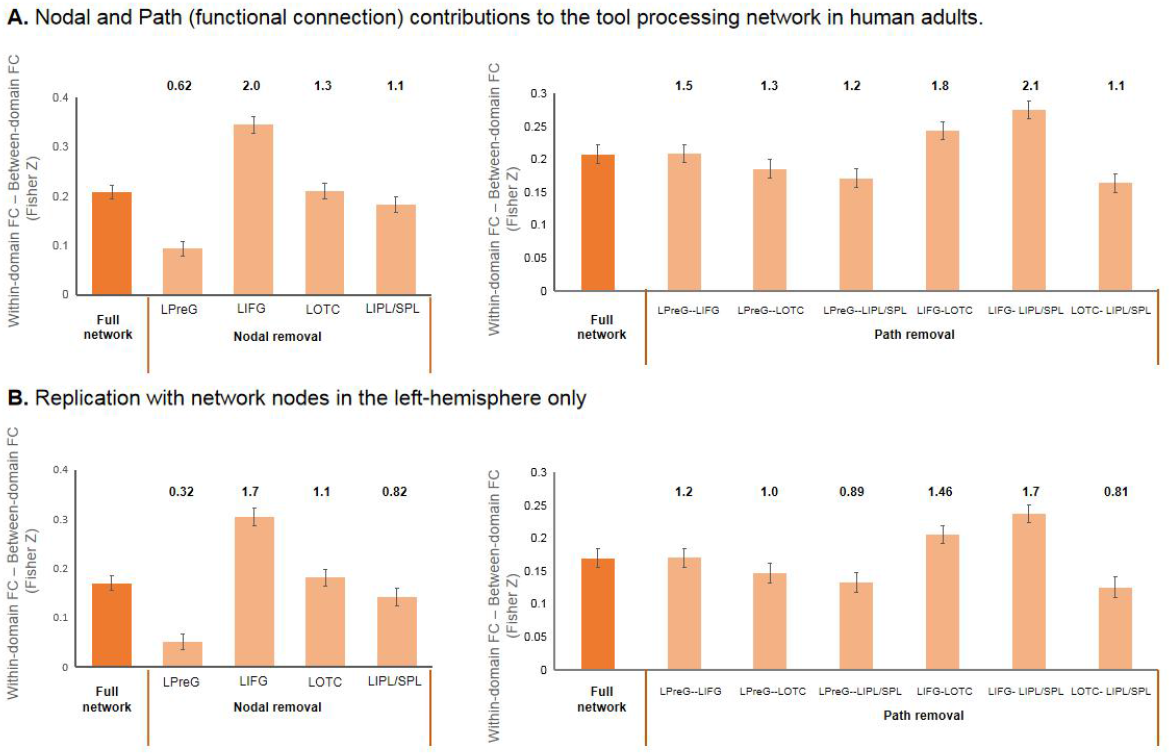
Nodal and path (functional connection) contributions to the tool processing network in human adults. A. Bar graphs illustrate network effects, calculated as within-domain minus between-domain rsFC, for the full tool processing network and when each constituent node (left column) or each constituent path (right column) is removed. Between-domain rsFC was computed from nodes in both hemispheres (Bilateral). Results show that no matter which node or path was removed, the within-domain rsFC strengths based on the remaining tool processing nodes or paths were still significantly higher than the between-domain rsFC strengths. Effect sizes (Cohen’s *d*) are shown above each bar. Error bars indicate standard errors. B. Bar graphs illustrate network effects, calculated as within-domain minus between-domain rsFC, for the full tool processing network and when each constituent node (left column) or each constituent path (right column) is removed. Between-domain rsFC was computed from nodes in left hemispheres. Results show that no matter which node or path was removed, the within-domain rsFC strengths based on the remaining tool processing nodes or paths were still significantly higher than the between-domain rsFC strengths. Effect sizes (Cohen’s *d*) are shown above each bar. Error bars indicate standard errors. LOTC: left lateral occipitotemporal cortex; LIPL/SPL: left inferior and superior parietal lobule; LPreG: left premotor gyrus; LIFG: left inferior frontal gyrus.

**Figure S6.**
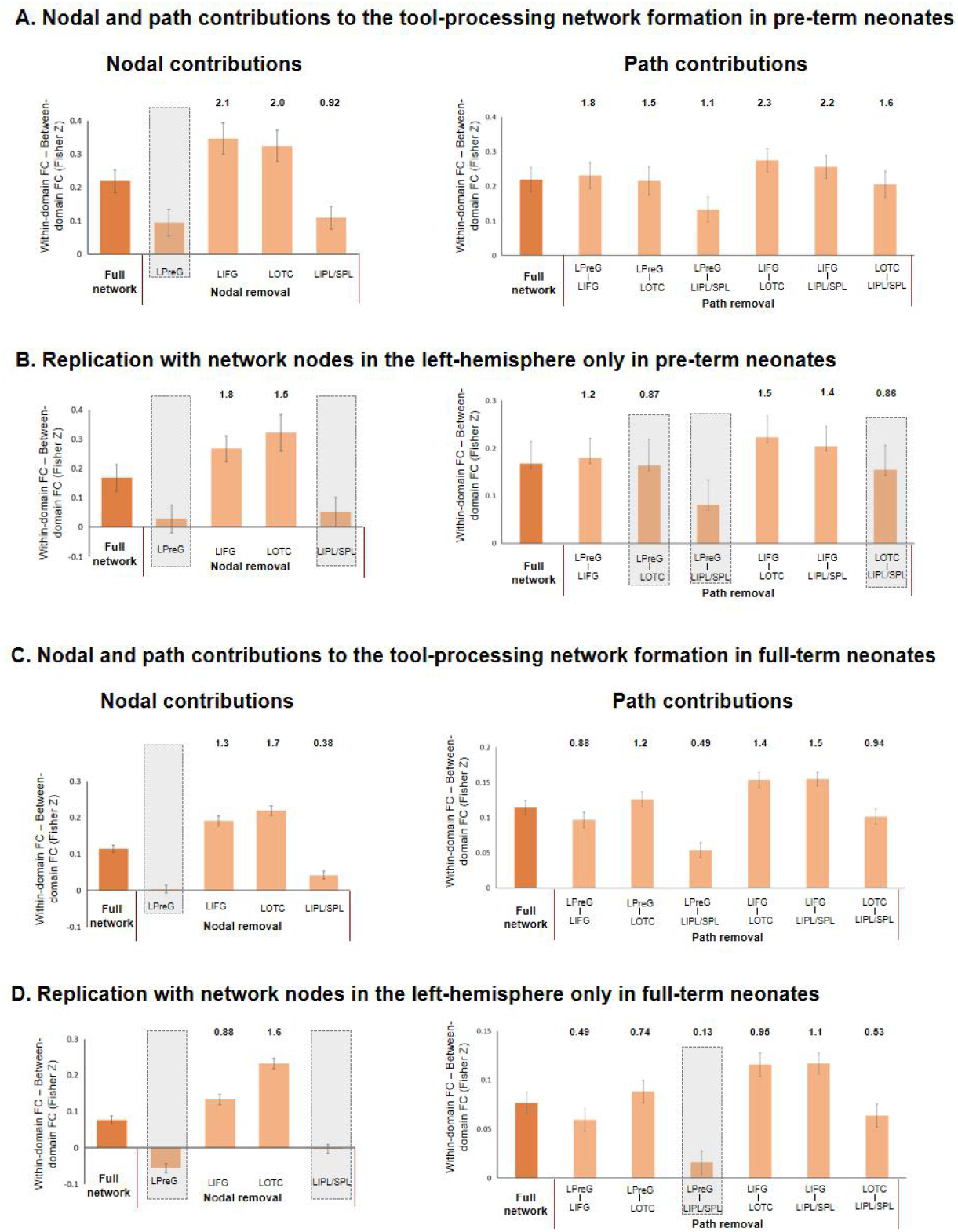
Critical contributions of the premotor node and its connectivity with the parietal region to the formation of the intrinsic tool processing network in pre-term and full-term neonate subgroups, as revealed by leave-one-node/path-out analyses. Bar graphs illustrate network effects, calculated as within-domain minus between-domain rsFC, for the full tool processing network and when each constituent node (left column) or path (right column) is removed. Between-domain rsFC was computed based on nodes in both hemispheres (bi-hemispheric, A for pre-term; C for full-term) or the left-hemisphere only (B for pre-term; D for full-term). Effect sizes (Cohen’s *d*) are shown when the rsFC strengths of the remaining tool network were still significantly higher than the between-domain rsFC strengths (all *p*_*corrected*_ < 0.05, Bonferroni corrected (nodal contribution: n= 4; path contribution: n = 6). Error bars indicate standard errors.

**Figure S7.**
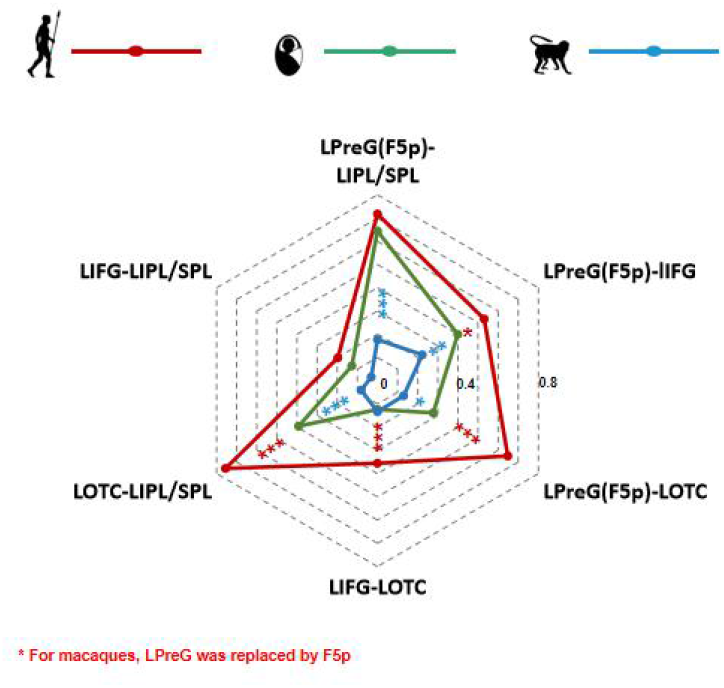
Path-specific connectivity strengths (Fisher-Z scores) of the tool (homologous) network in all three groups with LPreG replaced by F5p. Significant group comparisons between the human adults and human neonates are marked in *, whereas significant group differences between human neonates and macaques are marked in *. The left F5p (premotor)-left inferior/superior parietal path was species-specific, since it was the only path that was comparable between human adults and human neonates, but different between human neonates and macaques. LOTC: left lateral occipitotemporal cortex; LIPL/SPL: left inferior and superior parietal lobule; LPreG: left premotor gyrus; LIFG: left inferior frontal gyrus. * *p*_*corrected*_ < 0.05, ** *p*_*corrected*_ < 0.01, *** *p*_*corrected*_ < 0.001.

**Figure S8.**
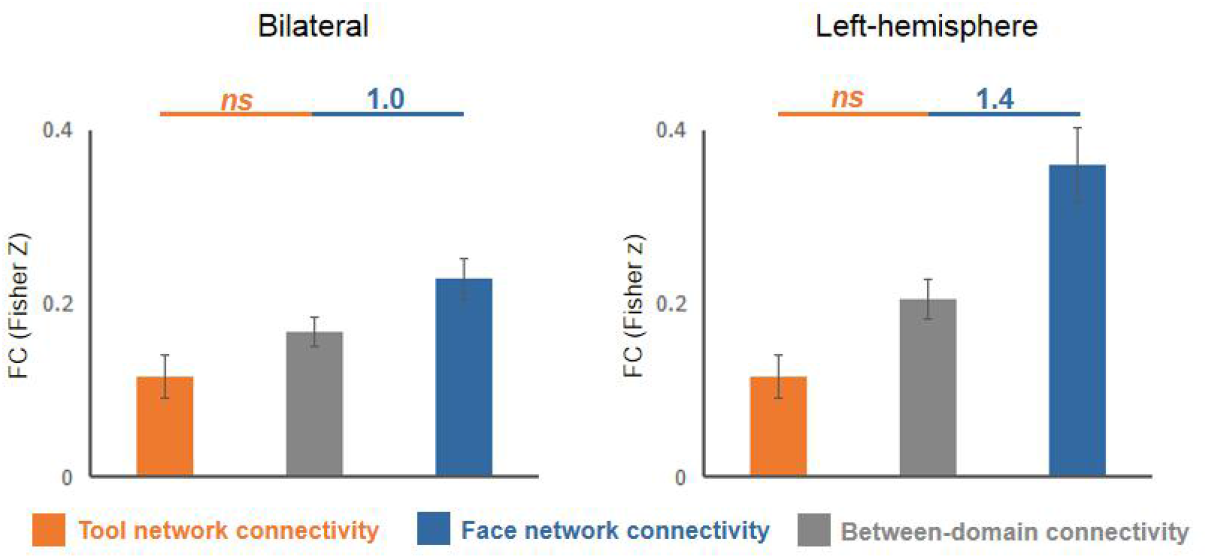
Replication results based on data from 17 macaques with at least 50% ROI coverage. Bar graphs illustrate rsFC values for within-domain and between-domain connectivity in the macaque group (with at least 50% ROI coverage) based on nodes in both hemispheres (bi-hemispheric, left column) or the left-hemisphere only (right column). Effect sizes (Cohen’s *d*) are shown for comparisons with significantly greater within-domain rsFC than between-domain rsFC (all *p*_corrected_ < 0.01, Bonferroni corrected (n = 2)). Error bars indicate standard errors.

